# Ganglioside GD3 regulates neural stem cell quiescence and controls postnatal neurogenesis

**DOI:** 10.1101/2023.03.14.532547

**Authors:** Takahiro Fuchigami, Yutaka Itokazu, Robert K. Yu

**Author notes:** Deceased May 18, 2022 (Robert K. Yu). Correspondence should be addressed to Y.I.: Yutaka Itokazu, Ph.D., Department of Neuroscience and Regenerative Medicine, Medical College of Georgia, Augusta University, 1120 15th Street Augusta, GA 30912, USA.

## Abstract

The postnatal neural stem cell (NSC) pool hosts quiescent and activated radial glia-like NSCs contributing to neurogenesis throughout adulthood. However, the underlying regulatory mechanism during the transition from quiescent NSCs to activated NSCs in the postnatal NSC niche is not fully understood. Lipid metabolism and lipid composition play important roles in regulating NSC fate determination. Biological lipid membranes define the individual cellular shape and help maintain cellular organization and are highly heterogenous in structure and there exist diverse microdomains (also known as lipid rafts), which are enriched with sugar molecules, such as glycosphingolipids. An often overlooked but key aspect is that the functional activities of proteins and genes are highly dependent upon their molecular environments. We previously reported that ganglioside GD3 is the predominant species in NSCs and that the reduced postnatal NSC pools are observed in global GD3-synthase knockout (GD3S-KO) mouse brains. The specific roles of GD3 in determining the stage and cell-lineage determination of NSCs remain unclear, since global GD3S-KO mice cannot distinguish if GD3 regulates postnatal neurogenesis or developmental impacts. Here we show that inducible GD3 deletion in postnatal radial glia-like NSCs promotes the NSC activation, resulting in the loss of the long-term maintenance of the adult NSC pools. The reduced neurogenesis in the subventricular zone (SVZ) and the dentate gyrus (DG) of GD3S-conditional-knockout mice led to impaired olfactory and memory functions. Thus, our results provide convincing evidence that postnatal GD3 maintains the quiescent state of radial glia-like NSCs in the adult NSC niche.

**Graphical abstract:** 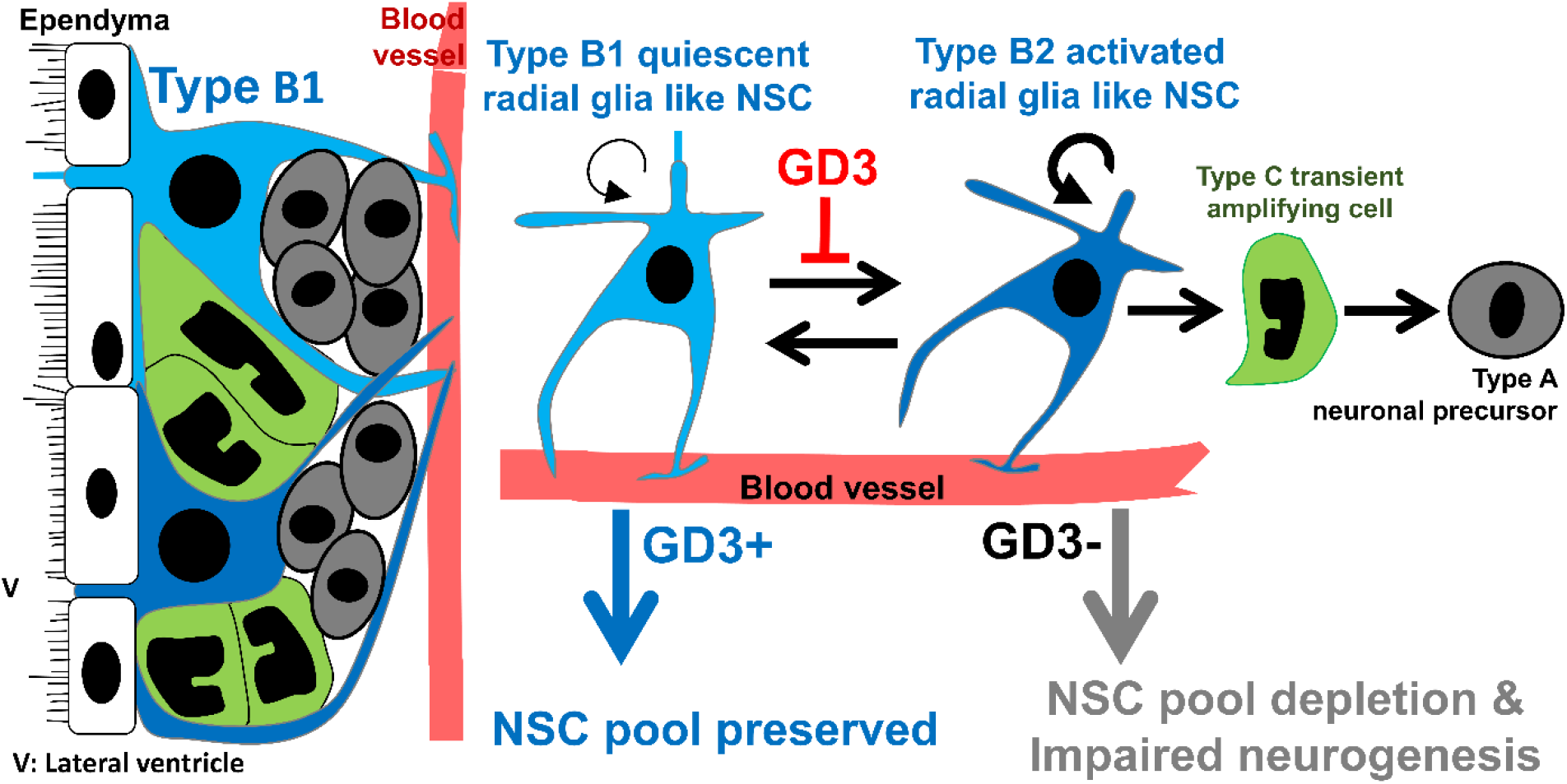

**Main Points:**

- Radial glia-like neural stem cells (RGLs) lacking GD3 promote activation of quiescent RGLs.
- Postnatal depletion of GD3 results in reduced neural stem cell pools and impaired adult neurogenesis.
- Postnatal GD3 deletion in RGLs leads to impairment of olfactory and memory functions.

## 1 INTRODUCTION

Postnatal neural stem cells (NSCs) mainly reside in the major neurogenic regions including the subventricular zone (SVZ) and the dentate gyrus (DG) of the hippocampus (Abbott & Nigussie, 2020; Codega et al., 2014; Doetsch et al., 1999; Ihrie & Alvarez-Buylla, 2011). Adult neurogenesis is a process of producing newborn neurons from radial glia-like NSCs (RGLs) in the adult mammalian brain throughout life. The NSC pool hosts quiescent and activated RGLs to sustain postnatal neurogenesis. Once quiescent RGLs are activated in the SVZ, they give rise to transient amplifying progenitors, which become DCX-expressing neural precursor cells (NPCs). NPCs migrate to the olfactory bulb (OB) through the rostral migratory stream (RMS). Migrated RGL-derived cells are located in the granule cell layer (GCL) or the periglomerular layer (PGL) of the OB where they eventually differentiate into interneurons (Ihrie & Alvarez-Buylla, 2011), which participate in the olfactory processing and odor discrimination (Sakamoto, Ieki, et al., 2014; Sakamoto, Kageyama, et al., 2014). The NSCs in the DG continuously supply intermediate progenitors, which would produce the granule neurons throughout life, which are involved in learning, memory, and general cognition (Abbott & Nigussie, 2020). An adult human generates 700 new neurons daily in the hippocampus, with a gradual decline during aging (Spalding et al., 2013). Therefore, the maintenance of the NSC pools and adult neurogenesis are suggested to be critical for keeping normal brain functions.

In the nervous system of vertebrates, lipids are the most abundant organic compounds, and a variety of lipids control the biophysical nature of lipid membranes. Biological membranes are highly heterogenous in structure and there exist diverse microdomains (also known as lipid rafts), which are enriched with glycosphingolipids (Itokazu & Yu, 2023). Glycosphingolipids are a class of lipids highly enriched in the nervous system, and are unique amphipathic molecules with a hydrophilic carbohydrate portion and hydrophobic lipid component (Yu et al., 2011). The significance is that glycosphingolipids modulate lipid microdomains to regulate functions of important molecules on plasma, mitochondrial, nuclear, and other biological membranes. Multifunctional gangliosides regulate specific cells in distinct stages via modulating protein and gene activities through the distinct ganglioside microdomains (Itokazu & Yu, 2023; Yu & Itokazu, 2014). Gangliosides are sialic acid-containing glycosphingolipids found in virtually all vertebrate cells, but primarily distributed on the outer cell membrane of the nervous system (Yu et al., 2009). Biosynthesis of gangliosides is catalyzed by glycosyltransferases (e.g., ganglioside synthase) in sequential steps. Abnormal regulation of ganglioside synthases and altered ganglioside expression has been associated with human development and diseases of nervous system (Itokazu et al., 2023). For example, loss of GM3 synthase leads to human autosomal recessive infantile-onset symptomatic epilepsy syndrome and Rett syndrome like disorder (Lee et al., 2016; Simpson et al., 2004). GM2 synthase deficiency results in hereditary spastic paralegias and axonal Charcot-Marie-Tooth syndrome (Boukhris et al., 2013; Hong et al., 2021; Lee et al., 2016; Simpson et al., 2004). Neurodegenerative diseases and mental health disorders are also associated with altered ganglioside expression (Itokazu et al., 2023). Accumulating studies demonstrate that gangliosides not only delimit physical regions but also play central roles in the maintenance of the biological functions of NSCs, neurons, and glia.

We first discovered that GD3 is the predominant ganglioside species, accounting for more than 80% of the total gangliosides in NSCs (Nakatani et al., 2010). GD3 is involved in the maintenance of NSCs in the SVZ and the DG of the postnatal brain by interacting with epidermal growth factor receptors on the cell surface and stabilizing its signal transduction. Further, the postnatal NSC pools are declined in the SVZ and the DG, and NPCs in the OB are reduced in global GD3-synthase-knockout (GD3S-KO) mice, suggesting impaired olfaction (Fuchigami et al., 2023; Itokazu et al., 2019; Itokazu et al., 2018; Tang et al., 2021; Wang et al., 2014; Wang & Yu, 2013). We have reported that the impaired neurogenesis in the adult of global GD3S-KO mice results in depressive symptoms and olfactory dysfunction and impaired memory. On the other hand, intranasal or intracerebroventricular infusion of GD3 restored postnatal NSC pools of global GD3S-KO mice (Fuchigami et al., 2023; Itokazu et al., 2019). These studies have shown that GD3 plays a pivotal role in maintaining the postnatal NSC pools to regulate neurogenesis and related behavior. Despite accumulating evidence obtained using global KO mice, the specific roles of GD3 in determining the stage and cell-lineage determination of NSCs remain unclear. Furthermore, global GD3S-KO mice cannot investigate if gangliosides regulate postnatal neurogenesis, since their gangliosides are eliminated before birth. The definitive test to show that the significance of postnatal GD3 in NSCs and neurogenesis would be to delete GD3 in postnatal NSCs.

To evaluate the role of GD3 in the postnatal NSCs, we generated GD3S floxed/floxed (f/f) mice with the mice carrying CreERT2 under Gli1 (glioma-associated oncogene homolog 1, GLI-Kruppel family member 1)-promoter with a stop-tdTomato reporter for inducible deletion of postnatal GD3. *In vivo* imaging and population-based lineage tracing have indicated that Gli1-CreERT2-labeled RGLs in the SVZ and the DG (Ahn & Joyner, 2005; Bottes et al., 2021; Guo et al., 2022) and those RGLs contribute to long-term maintenance of RGLs, self-renewal, and replenishment of interneuron subtypes in the OB (Ahn & Joyner, 2005; Bin Imtiaz & Jessberger, 2021; Bottes & Jessberger, 2021; Ihrie et al., 2011). In this study, we examined whether RGL-specific deletion of postnatal GD3 affects the NSC fate determination, e.g., quiescence, self-renewal, proliferation, differentiation, migration, and survival in the SVZ and the DG using Gli1-CreERT2;tdTomato; GD3Sf/f (GD3S-cKO) mice. By tracing the tdTomato (Tmt)-expressing cells, we identified newborn neurons after inducible deletion of GD3S. We found that RGL-specific GD3 deficiency led to reduction of the NSC pools and impaired neurogenesis in the SVZ and reduced Tmt+ cells in the OB, with impaired olfaction. Further, we discovered the decreased proliferative ability of RGL-derived cells and a declining trend of neurogenesis in the DG with impairment in hippocampus-dependent memory function. Taken together, the postnatal RGL-specific loss of GD3 resulted in reduced neurogenesis in both the SVZ and the DG, which led to functional impairment.

## 2 MATERIALS AND METHODS

### 2.1 Antibodies

For this study, the following antibodies were purchased: goat anti-tdTomato (RRID:AB_2722750, Origene, Rockville, MD, USA, #AB8181-200), rabbit anti-St8sia1 (RRID:AB_1857534; Atlas Antibodies, Stockholm, Sweden, #HPA026775), rabbit anti-Sox2 (RRID:AB_823640; Cell Signaling Technology, Danvers, MA, USA, #2748S), mouse anti-BrdU (RRID:AB_10015222; BD Biosciences, San Jose, CA, USA, #555627), mouse anti-Nestin (RRID:AB_396354; BD Biosciences, #556309), rabbit anti-glial fibrillary acidic protein (GFAP) antibody (RRID:AB_10013382; Dako Agilent, Santa Clara, CA, USA, #z0334), mouse anti-DCX (RRID:AB_10610966; Santa Cruz Biotechnology, CA, USA, #sc271390), mouse anti-proliferating cell nuclear antigen (PCNA) (RRID: AB_477413; Sigma, St. Louis, MO, USA, #P8825), rabbit anti-NeuN (RRID:AB_ 10807945; Millipore, St. Louis, MO, USA, #ABN78), rabbit anti-TH (RRID:AB_390204; Millipore, St. Louis, MO, USA, #AB152), rabbit anti-CR (RRID:AB_2228331; Synaptic Systems GmbH, Göttingen, Germany, #214 102), Alexa Fluor 488-conjugated goat anti-mouse immunoglobulin G (IgG) (RRID: AB_2536161; Invitrogen, #A28175), Alexa Fluor 488-conjugated goat anti-rabbit IgG (RRID:AB_143165; Invitrogen, #A11008), Alexa Fluor 555-conjugated donkey anti-goat IgG (RRID:AB_2535853; Invitrogen, #A21432), Alexa Fluor 568-conjugated goat anti-rabbit IgG (RRID:AB_143157; Invitrogen, #A11011), Alexa Fluor 647-conjugated goat anti-rabbit IgG (RRID:AB_2536101; Invitrogen, #A27040), Alexa Fluor 488-conjugated donkey anti-rabbit IgG (RRID:AB_2556546; Invitrogen, #R37118), and Alexa Fluor 488-conjugated donkey anti-mouse IgG (RRID: AB_2556542; Invitrogen, #R37114).

### 2.2 Animals

All animal experiments were approved by the Institutional Animal Care and Use Committee (IACUC) at Augusta University (AU) according to the National Institutes of Health (NIH) guidelines and were performed with approved animal protocols (references AUP 2009-0240 and 2014-0694). Gli1-CreERT2;tdTomato;GD3Sf/f (GD3S-cKO) mice were generated by crossing floxed allele of GD3S (GD3Sf/f) with Gli1-CreERT2 (The Jackson Laboratory, stock no. 007913) and TdTomato (Ai9; The Jackson Laboratory, stock no. 007909) mice. The GD3Sf/f mice were generated by injecting St8sia1^tm1a(EUCOMM)Hmg^ ES cells (reporter-tagged insertion with conditional potential) from International Mouse Strain Resource (EUCOMM, Munich, Germany) into the inner cell mass of C57BL/6J blastocysts. A floxed allele of the GD3S (St8sia1) gene contains loxP sites flanking Exon2. The injected blastocysts were then implanted into uterus of pseudo-pregnant foster mothers for further development. The LacZ-Neo cassette flanked by Flp recombinase recognition site (Frt) was removed by crossing mice carrying the GD3S-neo-lacZ-flox allele to FLPe transgenic mice (B6.Cg-Tg(ACTFlpE)9205Dym/J, The Jackson Laboratory, stock no 005703) which express a Flp recombinase to generate GD3S-loxP mice. The floxed mice were genotyped by PCR analysis of mouse tail DNA using primer set across the 3’ loxP site which are 5’-CAC CAC TGC CAT CAG AAC AC - 3’ and 5’-GGC TCT CCT CTC CCA TCT TC-3’. All the mice were backcrossed with C57BL/6J mice (The Jackson Laboratory, stock no. 000664) for more than 6–10 generations. The heterozygous male and female mice were mated and subjected to PCR screening for genotyping. Littermate GD3+/+ mice were used as controls. Tamoxifen (100 mg/ml, Cayman chemical, #132581, Ann Arbor, MI, USA) was freshly prepared in a 10% ethanol of corn oil (Sigma C8267), and injected to Gli1-CreERT2;tdTomato;GD3Sf/f mice daily for 5 consecutive days. BrdU (Sigma; 50 mg/kg body weight) was administered intraperitoneally to label newly dividing cells. All mice were housed in standard conditions with food and water provided *ad libitum* and maintained on a 12-hour dark/12-hour light cycle. Male mice were used in all experiments.

### 2.3 Immunohistochemistry

Mice were anesthetized with isoflurane using an open-drop method and transcardially perfused with phosphate-buffered saline (PBS, pH 7.4) and followed by 4 % paraformaldehyde (PFA). The brains were harvested and post-fixed with 4 % PFA overnight, followed by cryoprotection with 10% sucrose in PBS at 4°C overnight; then repeated more than three times with fresh 30% sucrose-PBS. After embedding in the Tissue-Tek OCT compound (Sakura Finetek, Torrance, CA, USA), the brains were quickly frozen in liquid nitrogen. Coronal or sagittal cryosectioning was performed at 20 μm using a cryostat (Leica, Wetzlar, Germany). For co-staining of tdTomato and GD3S, Sox2, BrdU, PCNA, Nestin, GFAP, DCX, or NeuN, sections were treated with microwave for 5 minutes in pre-boiled 10 mM citrate buffer (pH 6.0), followed by permeabilization with PBS containing 0.5 % Triton X-100 for 5 min and blocked with PBS containing 1 % bovine serum albumin (BSA) for 30 min at room temperature, and then incubated with goat anti-tdTomato antibody (1:300, Origene, # AB8181-200), rabbit anti-St8sia1 (GD3S) (1:100, Atlas Antibodies, #HPA026775), anti-PCNA antibody (1:100, Sigma, #P8825), mouse anti-DCX antibody (1:50, Santa Cruz, #sc271390), rabbit anti-NeuN antibody (1:100, Millipore, #ABN78), rabbit anti-Sox2 antibody (1:100, Cell Signaling Technology, #27485) and mouse anti-BrdU antibody (1:500, BD Biosciences, #555627) or mouse anti-Nestin antibody (1:100, BD Biosciences, #556309) and rabbit anti-GFAP antibody (1:100, Dako Agilent, #z0334) at 4°C overnight. Nuclei counterstaining was performed with 1 μg/mL 40,6-diamidino-2-phenylindole (DAPI) (Thermo Fisher Scientific, #D1306) for 30 min. After every incubation with antibodies or chemicals, sections were washed three times with PBS. Specimens were mounted with VectaMount (Vector Laboratories, Burlingame, CA, USA).

### 2.4 Microscopy and image processing

Images of labeled sections were acquired by a Zeiss LSM700 (Carl Zeiss, Land Baden-Württemberg, Germany) with a 63x (1.4 NA oil immersion, Plan Apochromat) objective as 10 μm of z-stacks of optical slices or a Nikon A1R MP+ multiphoton/confocal microscope (Nikon, Tokyo, Japan) with an ApoLWD 25 x (1.10 W DIC, N2) objective. The microscope settings were kept constant for each staining. Serial z-images were stacked using Fiji (NIH, Bethesda, MD, USA) and merged using Photoshop (Adobe, San Jose, CA, USA). tdTomato, Nestin, GFAP, Sox2, DCX, PCNA, and BrdU in the SVZ were counted at the DAPI-stained germinal zone for quantification. NeuN-, tyrosine hydroxylase (TH)-, and calretinin (CR)-labeled DAPI-stained cells were counted in the PGL of the OB for each group (n= 5 mice, 5 sections per each mouse). Immunofluorescence higher than background and specific staining pattern/localization were considered to determine positive/negative for each staining when cells were counted using Fiji image analysis. To calculate the total number of marker-positive cells, 5 sections (200 μm apart) per animal and at least 1,000 cells per group were analyzed. The blinding procedures and randomized field approach for images were performed to acquire unbiased results.

### 2.5 Buried pellet test

The control (GD3S+/+) and GD3S-cKO mice (4-month-old mice; male) were fasted for 24h before testing in a home cage with water supply. On the day of the experiment, the mice were habituated in a testing cage with 3 cm depth bedding for 10 min and placed back in the home cage. Then a food pellet (1 cm^3^) was placed under the bedding in the testing cage. The mouse was placed in the center of the testing cage, and the latency (seconds) to uncover the food pellet was measured. If the mouse did not find the pellet within 5 min, the trial was finished scoring 300 sec for the mouse. The time until mice uncovered the food pellet in bedding was measured as an average of 4 trials. In each trial, a pellet was embedded beneath the bedding at a different position. After all the mice were tested, they were fed. Behavioral experiments were performed between 8 and 11 am.

### 2.6 Novel object recognition test

After 10 min of habituation in the box (40 cm by 40 cm by 30 cm), mice were placed in the center of the arena with two identical objects in a symmetrical position of 5 cm from the walls. The mice were returned to the home cage after the training session. 1h and 24h after the training, mice were tested for novel object recognition for 5 min each, in which one of the objects was replaced by a new object with a different shape and color. The mice were placed at the center of the chamber at the beginning of every trial. The chamber and the objects were wiped with 70% ethanol prior to every trial to avoid odor effects. Exploration of the objects was defined as directing the nose or sniffing, touching, and sitting on it as attracted towards the objects while the time just moving around the objects was not recorded. The activity was digitally recorded with Ethovision XT video-tracking software (Noldus Information Technology, Asheville, NC).

### 2.7 Statistical analysis

All statistical procedures were performed using GraphPad Prism 9 (GraphPad, San Diego, CA, USA). Group data were analyzed using the Kolmogorov-Smirnov test to determine their normality, and F-test examined the homogeneity of variances of datasets. An unpaired two-tailed Student’s *t*-test was performed when applicable. If not, Mann Whitney test was used to compare differences. In all cases, *p* values < 0.05 were regarded as significant, as shown in the figure legends. All data were depicted as the mean ± SEM. Our sample sizes were determined by previous study (Fuchigami et al., 2023; Tang et al., 2021). To estimate the biological variability, ensure the quality and consistency between biological samples and meanwhile maximize the cost-effectiveness, we chose to use 5–12 animals per condition. In box and whisker plot (Figures 6 and 8), a box range represents upper and lower quartiles, and the end of whiskers represent the minimum and maximum values outside of the upper and lower quartiles. The median values are represented by bars in the boxes.

**Figure 1.**
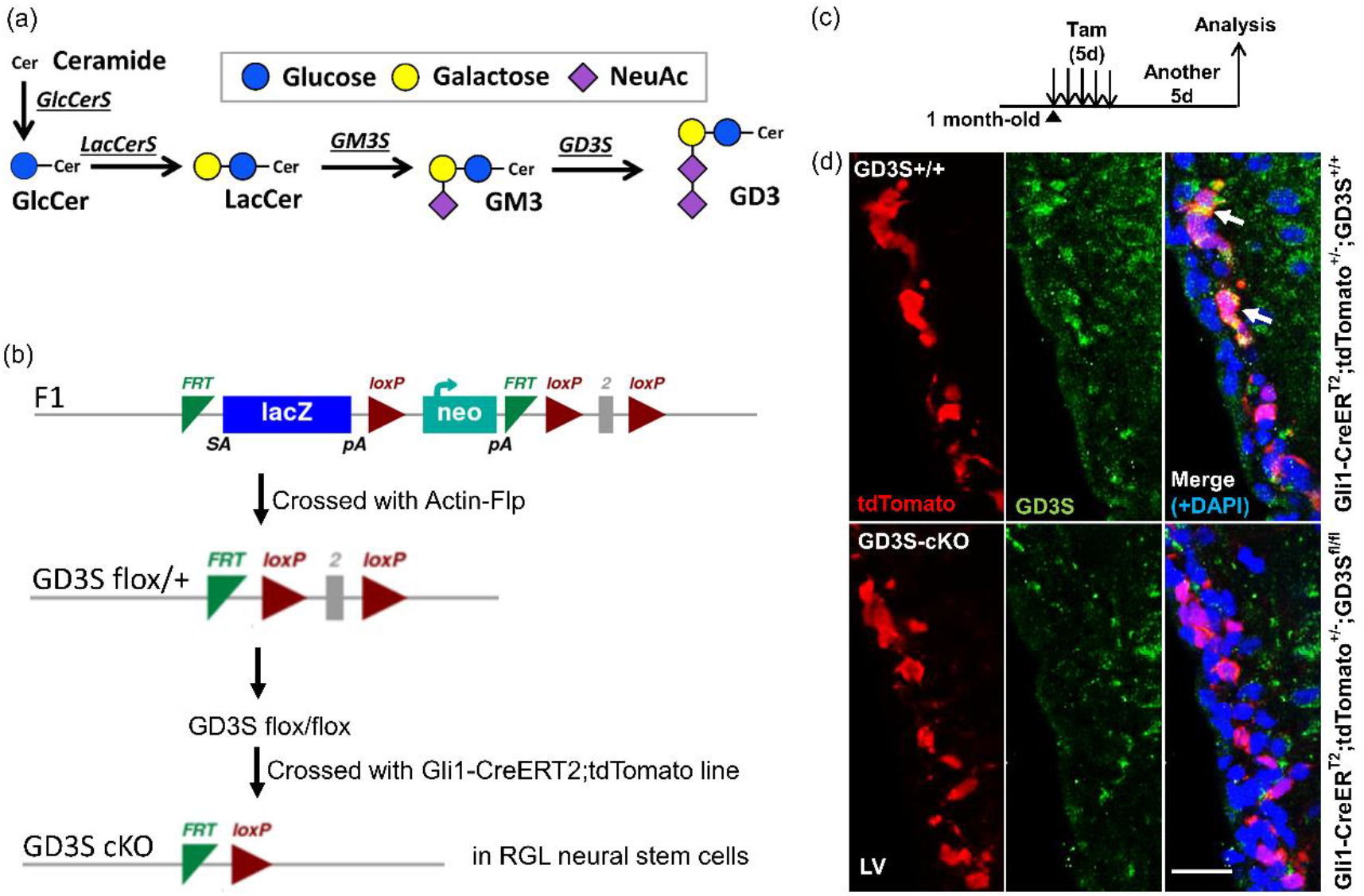
Generation of radial glia-like neural stem cell specific GD3S-conditional knockout mice. (a) Metabolic pathways and structure of ganglioside GD3. Glycosyltransferases (underlined) catalyze the biosynthesis of gangliosides. GD3S is a critical enzyme for the synthesis of GD3 from GM3. GlcCerS: Glucosylceramide synthase, Ugcg; LacCerS: Lactosylceramide synthase, B4galt5; GM3S: GM3 synthase, St3gal5, GD3S: GD3 synthase, St8Sia1; NeuAc: N-Acetylneuraminic Acid. (b) Schematic of generation of radial glia-like neural stem cell (RGL) specific GD3S-conditional knockout, Gli1-CreERT2;GD3Sf/f (GD3S-cKO) mice. First panel illustrates that the L1L2_Bact_P cassette was inserted at position 142914589 of chromosome 6 upstream of the Exon2 of GD3S in the ES cell line, The cassette is composed of an FRT site followed by lacZ sequence and a loxP site. This first loxP site is followed by a neomycin resistance gene under the control of the human beta-actin promoter, SV40 polyA, a second FRT site and a second loxP site. A third loxP site was inserted downstream of the Exon2 at position 142913744. The exon2 of GD3S gene is thus flanked by loxP sites. In the second step, the floxed allele was created by crossing the F1 mice with flp recombinase expressing mice (Actin-Flp). Subsequent cre expression resulted in the generation of knockout mice. At the third step, we generated GD3S-cKO mice. Gli1-CreERT2;tdTomato;GD3Sf/f (GD3S-cKO) mice were generated by crossing floxed allele of GD3S (GD3Sf/f) with Gli1-CreERT2 and TdTomato (Ai9) mice. (c) Summary of the experimental design. 1-month-old mice were treated with Tamoxifen for 5 days and followed by analysis 5 more days later. (d) Immunohistochemistry for tdTomato (Tmt, red) and GD3S (green) combined with DAPI (blue) on coronal 20 μm cryosections. GD3S signal was depleted in the SVZ of cKO mouse brain. LV: lateral ventricle. Scale bar, 20 μm.

**Figure 2.**
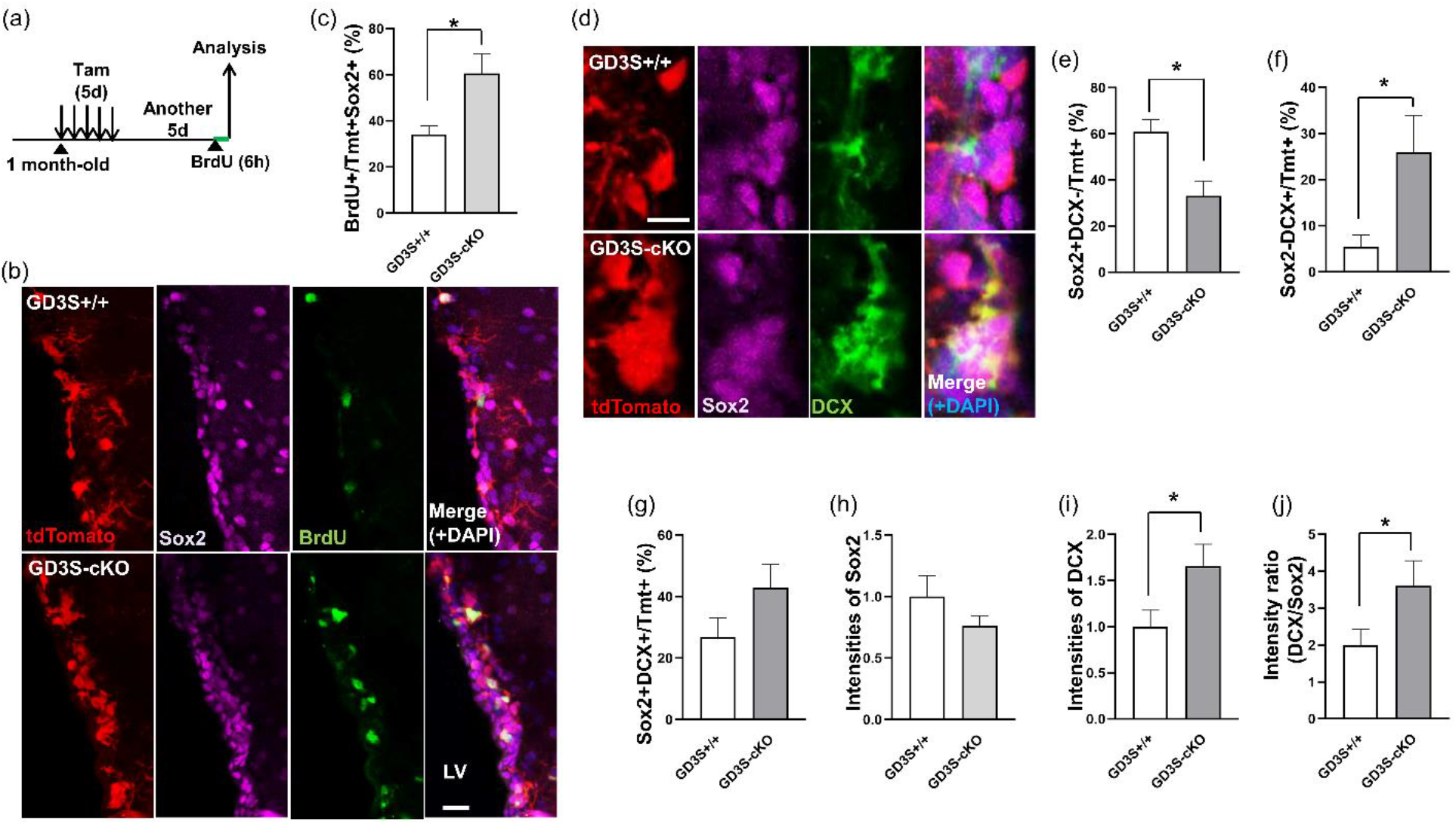
GD3-deficient RGLs markedly decreased quiescence and promoted to proliferate and differentiate to the neural precursor cells in the SVZ of GD3S-cKO mice at 5 days after tamoxifen injection. (a) Summary of the experimental design. 1-month-old mice were treated with Tamoxifen for 5 days and followed by analysis 5 more days later. (b) Immunohistochemistry for Tmt (red), Sox2 (magenta), and BrdU (green) combined with DAPI (blue) on coronal 20 μm cryosections. Scale bar, 20 μm. LV: lateral ventricle. (c) Quantification of BrdU-incorporated Tmt+Sox2+ cells after recombination and its statistical analysis. *; p<0.05. (d) Immunohistochemistry for Tmt (red), Sox2 (magenta), and DCX (green) combined with DAPI (blue) on coronal 20 μm cryosections. Scale bar, 10 μm. (e–g) Quantification of (e) Tmt+Sox2+DCX-, (f) Tmt+Sox2-DCX+, and (g) Tmt+Sox2+DCX+ cells and their statistical analysis. *; p<0.05. (h, i) Quantitative analysis for intensities of Sox2 (h) and DCX (i) within Tmt+ cells. Mean ± SEM. *; p<0.05. (j) Intensity ratio of DCX to Sox2. Mean ± SEM. *; p<0.05. n= 5 mice/group.

**Figure 3.**
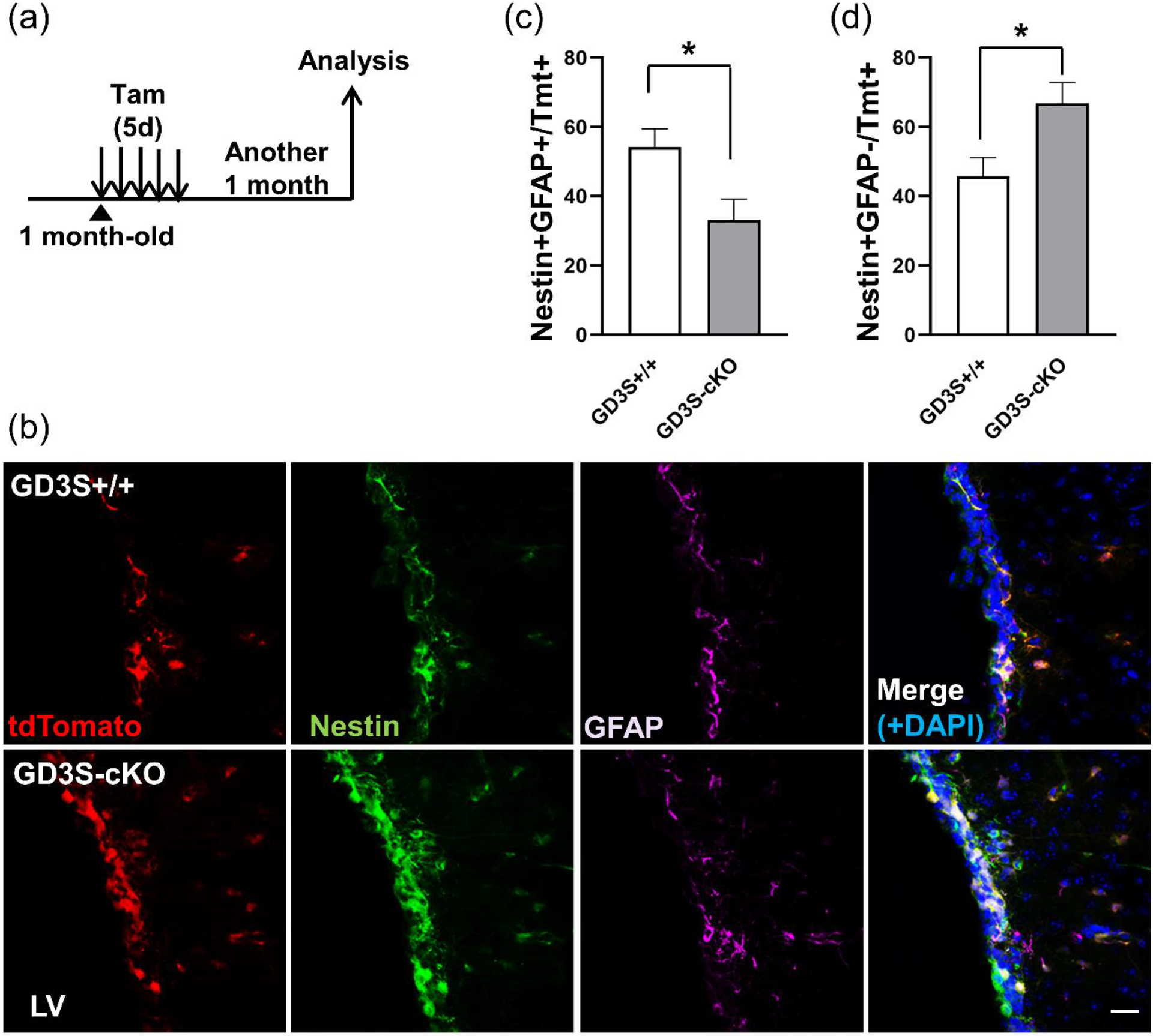
Nestin+GFAP+ radial glia-like quiescent neural stem cells decreased in GD3-deficient RGL at the SVZ of GD3S-cKO mice at 1 month after tamoxifen injection. (a) Summary of the experimental design. 1-month-old mice were treated with Tamoxifen for 5 days and followed by analysis 1 more month later. (b) Immunohistochemistry for Tmt (red), Nestin (green), and GFAP (magenta) combined with DAPI (blue) on coronal 20 μm cryosections. Scale bar, 20 μm. LV: lateral ventricle. Quantification of (c) Nestin+GFAP+ and (d) Nestin+GFAP-in RGL-derived cells (Tmt+) from imaging data of (b). *; p<0.05. n= 5 mice/group.

**Figure 4.**
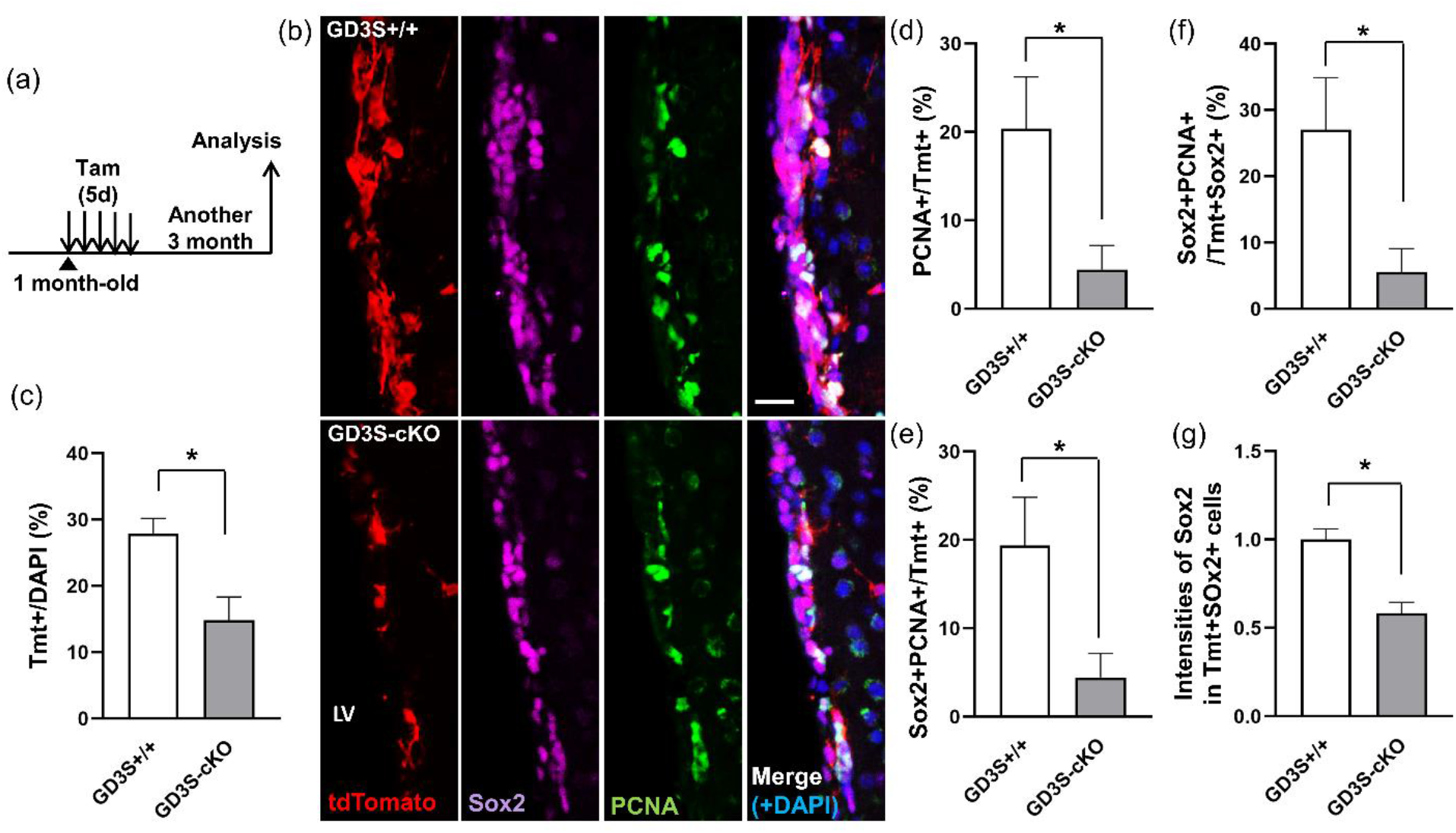
GD3 deletion in radial glia-like neural stem cells (RGLs) reduced the NSC pool in the SVZ of the GD3S-cKO mice at 3 month after tamoxifen injection. (a) Summary of the experimental design. 1-month-old mice were treated with Tamoxifen for 5 days and followed by analysis after 3 more months. (b) Immunohistochemistry for Tmt (red), Sox2 (magenta), and PCNA (green) combined with DAPI staining (blue) on coronal 20 μm cryosections. Scale bar, 20 μm. LV: lateral ventricle. (c–f) Quantification of (c) Tmt+ RGL-derived cells, (d) Tmt+PCNA+ dividing cells among Tmt+ cells, (e) Tmt+Sox2+PCNA+ activated NSCs among Tmt+ RGL-derived cells, and (f) Tmt+Sox2+PCNA+ activated NSCs in Tmt+Sox2+ RGL-derived NSCs. *; p<0.05. (g) Quantification of the intensity of Sox2 within Tmt+ cells. Mean ± SEM. *; p<0.05. n= 5 mice/group.

**Figure 5.**
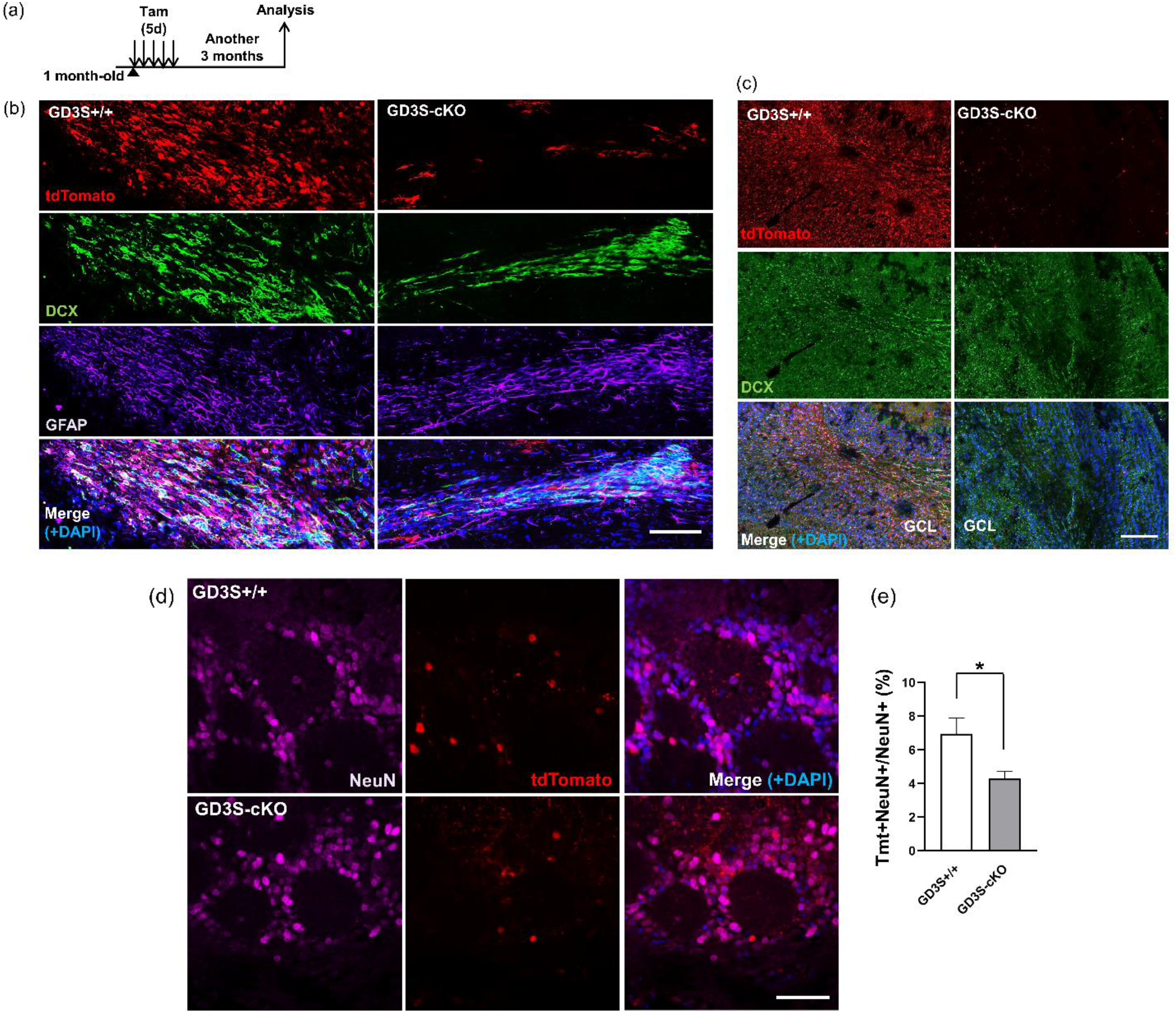
Reduced neurogenesis from GD3-deficient radial glia-like neural stem cells (RGLs) was found in the RMS-OB of GD3S-cKO mice. (a) Summary of the experimental design. 1-month-old mice were treated with Tamoxifen for 5 days and followed by analysis 3 more months later. (b) Immunohistochemistry for Tmt (red), DCX (green), GFAP (magenta) combined with DAPI staining (blue) on sagittal 20 μm cryosections (RMS: rostral migratory stream). Scale bar, 100 μm. (c) Immunohistochemistry for Tmt (red) and DCX (green) combined with DAPI (blue) in the granule cell layer (GCL) of the OB. Scale bar, 200 μm. (d) Immunohistochemistry for Tmt (red) and NeuN (magenta) combined with DAPI (blue) in the periglomerular layer (PGL) of the OB. Scale bar, 50 μm. (e) Quantification of Tmt+ cells among NeuN+ cells from imaging data of (d). Tmt+NeuN+ cells were decreased in GD3S-cKO mice.. *; p < 0.05. n= 5 mice/group.

**Figure 6.**
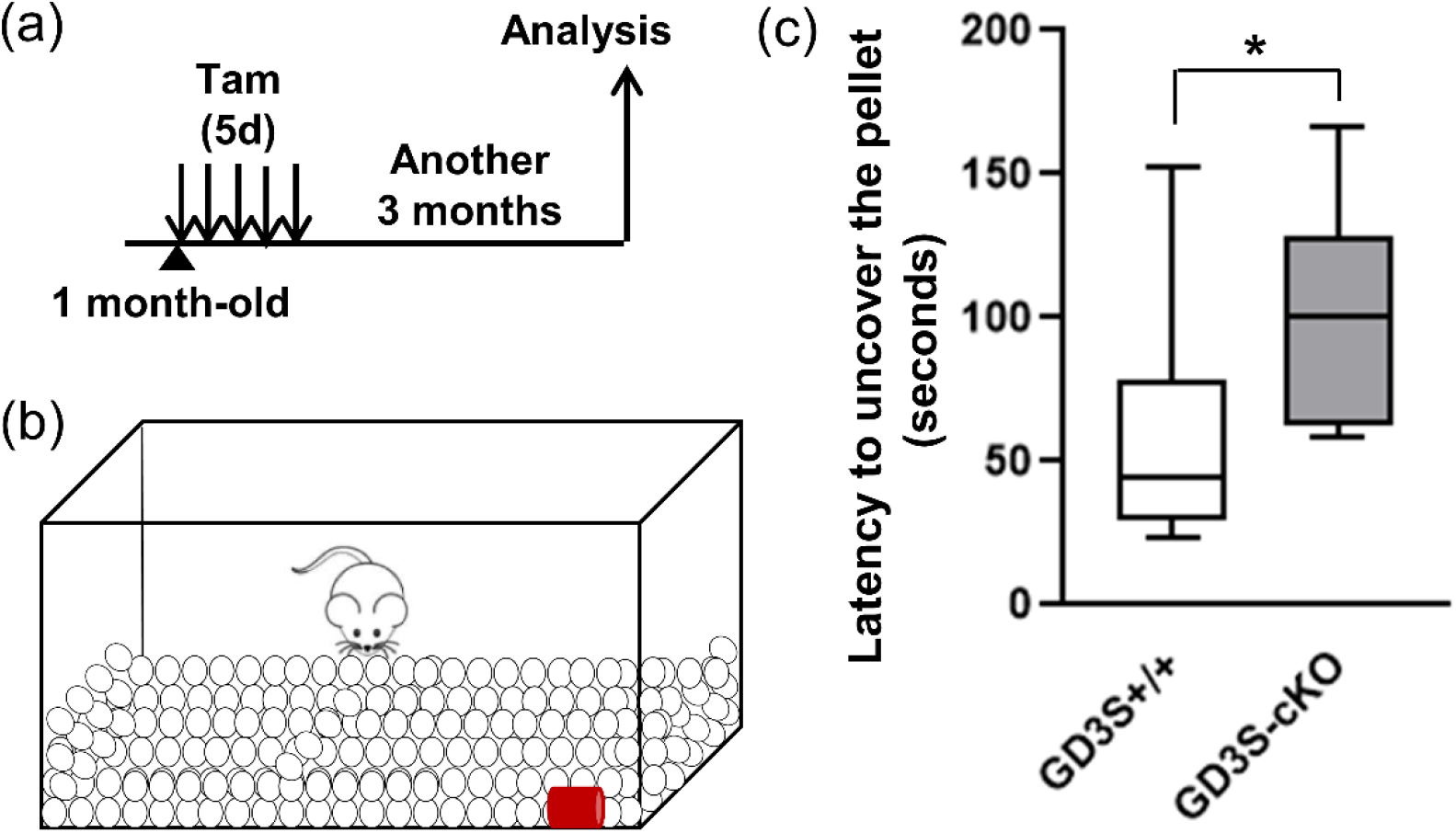
Induced ablation of postnatal GD3 in radial glia-like neural stem cells (RGLs) impairs olfaction in GD3-cKO mice. (a) Summary of the experimental design. 1-month-old mice were treated with Tamoxifen for 5 days and followed by analysis 3 more months later. (b) Schematic diagram showing the buried pellet test. The buried pellet test, which relies on the animal’s natural tendency to use olfactory cues for foraging, is used to confirm the ability to smell. The time until mice uncovered the food pellet in bedding was measured as an average of 4 trials. (c) Quantification of the latency (seconds) to uncover the buried pellet. The box range represents the upper and lower quartiles, and the end of the whiskers represents the minimum and maximum values. The median values are represented by bars in the boxes. *; p < 0.05. n= 12 for the control and n= 7 for GD3S-cKO.

**Figure 7.**
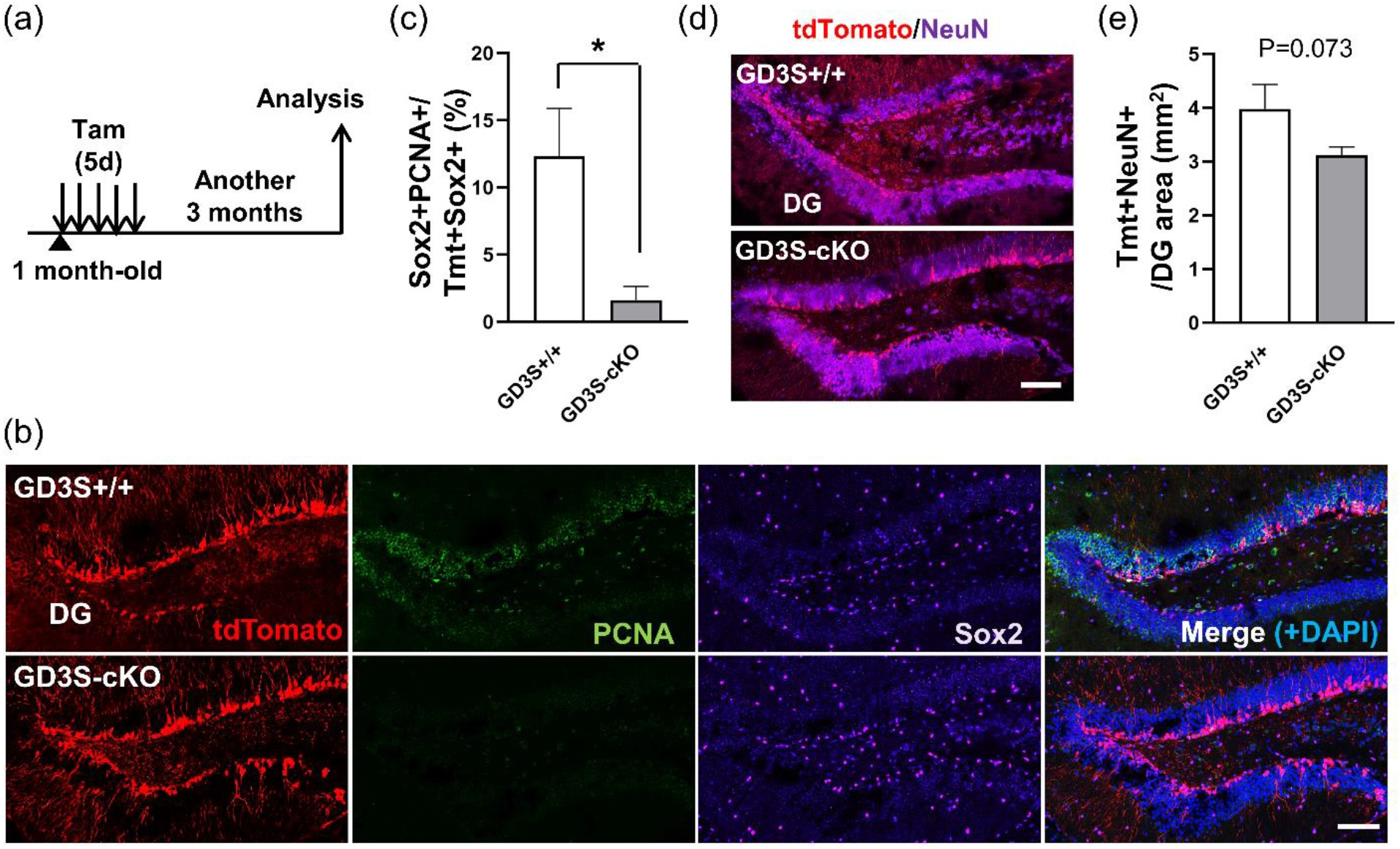
Reduced neurogenesis from GD3-deficient radial glia-like neural stem cells (RGLs) was found in the DG of GD3S-cKO mice. (a) Summary of the experimental design. 1-month-old mice were treated with Tamoxifen for 5 days and followed by analysis 3 more months later. (b) Immunohistochemistry for Tmt (red), PCNA (green), and Sox2 (magenta) combined with DAPI (blue) in the dentate gyrus (DG). Scale bar, 100 μm. (c) Quantification of Sox2+PCNA+ proliferating-active NSCs among Tmt+ RGL-derived cells. *; p < 0.05. (d) Immunohistochemistry for Tmt (red) and NeuN (magenta) combined with DAPI (blue) in the DG. Scale bar, 100 μm. (e) Quantification of Tmt+NeuN+ newborn neurons in the DG area. *; p < 0.05. n= 5 mice/group.

**Figure 8.**
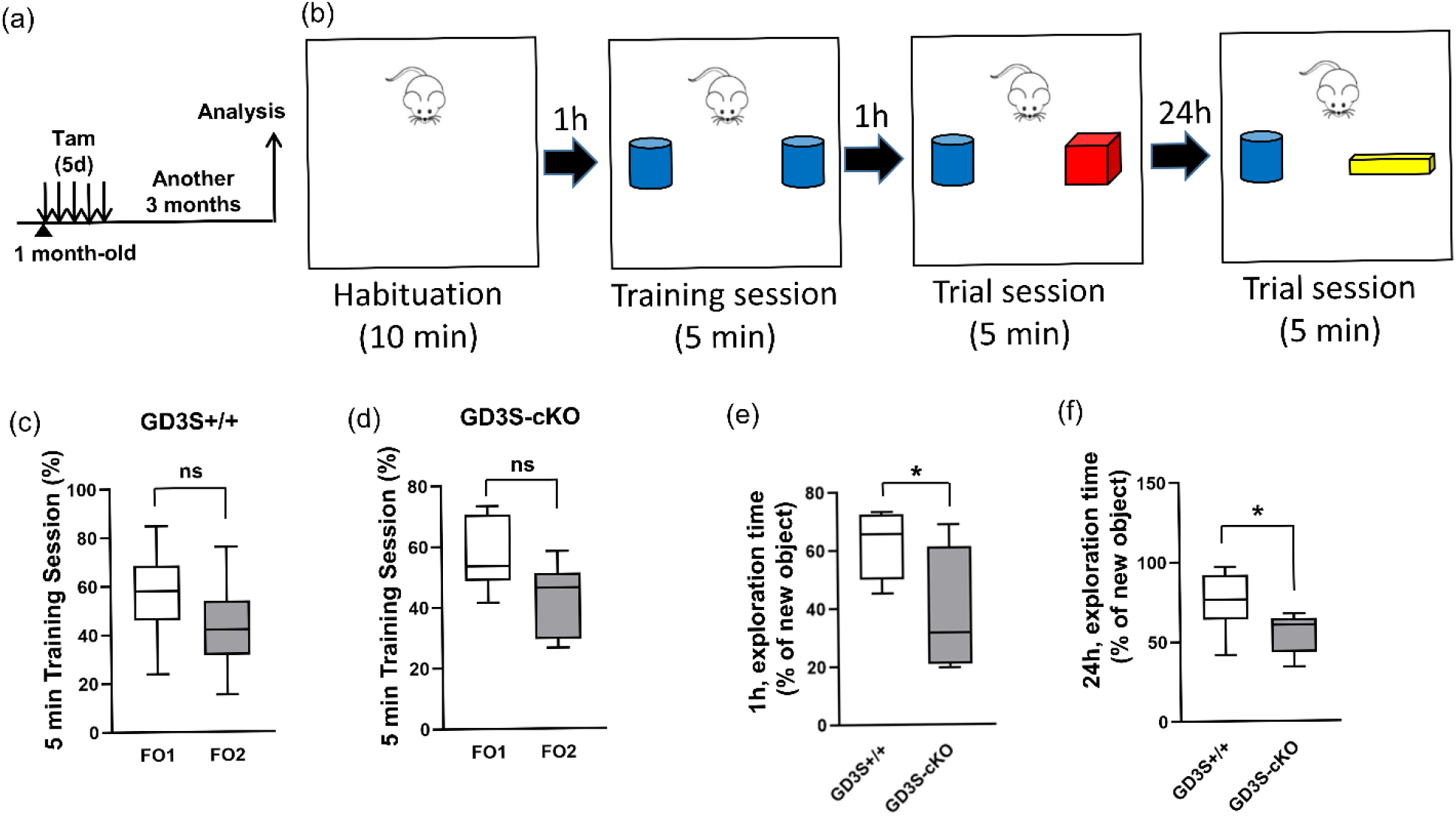
Induced ablation of postnatal GD3 in radial glia-like neural stem cells (RGLs) impairs hippocampus-dependent memory in GD3-cKO mice. (a) Summary of the experimental design. 1-month-old mice were treated with Tamoxifen for 5 days and followed by analysis 3 more months later. (b) Experimental procedure of the novel object recognition test. (c, d) The proportion of the exploring time for two familiar objects in the training session. (e, f) The proportion of the exploring time for the novel object to the familiar object 1h (e) or 24h (f) after the training. The box range represents the upper and lower quartiles, and the end of the whiskers represents the minimum and maximum values. The median values are represented by bars in the boxes. *; p < 0.05. n=12 for the control and n= 6 for GD3S-cKO.

## 3 RESULTS

### 3.1 Inducible GD3 deletion in RGLs results in activation

It has been revealed that GD3 is required for long-term maintenance of NSC populations in the postnatal mouse brain. Deficiency in GD3 leads to developmental and behavior deficits, such as depression, impairment in hippocampus-dependent memory, and olfactory dysfunction. GD3S is the critical enzyme for the synthesis of GD3 from GM3 (Figure 1a). Albeit amount of evidence by utilizing global GD3S-KO mouse, age- and cell lineage-specific role of this ganglioside in the postnatal brain remains unknown. It is conceivable that reduced adult neurogenesis in global GD3S-KO animal may be due to loss of GD3 in embryonic stage and/or due to lack of GD3 function in NSCs at adult stage. The inducible Cre-loxP system provides a selective deletion of genes that are essential for proper ganglioside synthesis and enables the study of ganglioside functions in postnatal NSCs. To establish the GD3S floxed mice, we made chimeric mice with an ES cell line with GD3S-targeted mutation (reporter tagged insertion with conditional potential) from International Mouse Strain Resource. We obtained 4 chimeric mice and have germline transmission of F1. The F1 mice were crossed with transgenic C57BL/6J mice expressing an enhanced variant of FLP1 recombinase, under the actin promoter. The offspring were backcrossed onto C57BL/6J background for >6 generations (Figure 1b).

We asked what happens when we inducibly delete GD3S in postnatal RGLs. To assess the role of GD3 in the postnatal NSCs, we bred GD3Sf/f mice with the mice carrying CreERT2 under Gli1-promoter with a stop-tdTomato reporter. We chose the Gli1-CreERT2 driver line, since Gli1-CreERT2-labeled RGLs in the SVZ and the DG (Ahn & Joyner, 2005; Bottes et al., 2021) and those RGLs commit to longstanding maintenance of NSCs and interneuron neurogenesis in the OB (Ahn & Joyner, 2005; Bin Imtiaz & Jessberger, 2021; Bottes & Jessberger, 2021; Ihrie et al., 2011). Administration of tamoxifen (5 consecutive daily) induced deletion of GD3S and tdTomato (Tmt) expression in the SVZ of GD3S-cKO mice, while cytosolic distribution of GD3S was found in Tmt+ cells in the SVZ of the control mice (Figure 1c,d).

We induced GD3S recombination and Tmt expression in RGLs of postnatal 1 month old, and injected BrdU to label dividing cells for 6 h prior to sacrifice (Figure 2a). We found that conditional deletion of GD3S in Gli1-CreERT2-targeted RGLs significantly increased the fraction of proliferating active RGLs. BrdU-incorporated Tmt+Sox2+ cells were increased in the SVZ of GD3S-cKO mice (control; 33.3 ± 3.53% vs. GD3S-cKO; 57.6 ± 8.48%) (Figure 2b,c). Figure 2b shows that radial glial processes were impaired in GD3S-cKO cells, while GD3+ RGLs had more complex morphologies of radial glia like morphology. Since it is known that RGLs divide symmetrically to self-renew or asymmetrically to generate an RGL and a DCX-expressing neuronal precursor cell (Ihrie & Alvarez-Buylla, 2011; Kaneko et al., 2017; Yang et al., 2004), we examined the expressions of Sox2 and DCX in RGL-derived Tmt expressing cells in the SVZ of the control and GD3S-cKO mice (Figure 2d). We found a decreased number of Tmt+Sox2+DCX-NSCs (control; 60.8 ± 5.46% vs. GD3S-cKO; 33.2 ± 6.36%) (Figure 2e) and an increased number of Tmt+Sox2-DCX+ neuronal precursor cells (control; 5.5 ± 2.48% vs. GD3S-cKO; 26.0 ± 7.94%) in GD3S-cKO mice (Figure 2f). Tmt+Sox2+DCX+ cells appeared to be increased, which might be becoming to Sox2-DCX+ neuronal precursor cells (control; 26.8 ± 6.37% vs. GD3S-cKO; 43.1 ± 7.47%) (Figure 2g). Next, we quantified the expression levels of Sox2 and DCX by measuring their fluorescent intensities to examine the cellular differentiation state in the SVZ. As expected, GD3S-cKO mice exhibited higher DCX levels (1.66 ± 0.238-fold vs. control) without significant difference in the Sox2 expression (Figure 2h,i). Consistently, the fluorescent intensity ratio of DCX to Sox2 was higher in the GD3S-cKO mouse (1.81 ± 0.341-fold vs. control) (Figure 2j). Together, these results demonstrate that GD3 is enriched in quiescent RGLs and that loss of GD3 expression in RGLs promotes an activated state of RGLs and differentiation. These results thus support the idea that chronic deletion of GD3 would result in a shrink of the NSC pool in the SVZ.

### 3.2 Inducible GD3 deletion in RGLs results in loss of stemness

To investigate the longer-term effect of GD3S depletion, tamoxifen was administered at 1-month-old and we analyzed RGL-derived lineage at 1 month after tamoxifen injection (2-month-old) (Figure 3a). The results showed a reduced number of Nestin+ GFAP+ RGLs in the SVZ of GD3S-cKO mice compared to the controls (control; 54.1 ± 5.34% vs. GD3S-cKO; 33.2 ± 5.99%, p = 0.04) (Figure 3b,c). Consistently, the number of Nestin+GFAP-cells was increased in GD3S-cKO mice (control; 45.8 ± 5.34% vs. GD3S-cKO; vs. 66.8 ± 5.99%, p = 0.04) (Figure 3d). To further clarify the effect of postnatal GD3 on long-term neurogenesis, we analyzed RGL-derived lineage at 3 months after tamoxifen injection (4-month-old) (Figure 4a). We found that Tmt+ cells were significantly decreased in the SVZ of cKO compared to the control mice (control; 27.9 ± 2.31% vs. 14.9 ± 3.50 %) (Figure 4b,c). Continuous neurogenesis throughout life depends on the pool of NSCs that self-renew, proliferate, and differentiate into multiple neural cell types. To evaluate the proliferating cell population of the RGL-derived cells, immunohistochemistry was performed to quantify PCNA expressing cells. PCNA+ proliferating cells in the RGL-derived cells (Tmt+, control; 20.4 ± 5.85% vs. GD3S-cKO; 4.44 ± 2.72%) (Figure 4d). Proliferating Sox2+ cells were also decreased among the Tmt+ cells (control; 19.4 ± 5.48% vs. GD3S-cKO; 4.44 ± 2.72%) (Figure 4e) and the Sox2+ Tmt+ cells (control; 27.0 ± 7.85% vs. GD3S-cKO; 5.56 ± 3.51%) (Figure 4f). Although analysis of NPCs’ proliferation based on the cell cycle marker PCNA cannot distinguish the actively dividing NSCs or the transient-amplifying progenitors, the reduction of proliferating cells implies a progressive loss of neurogenesis capacity in the GD3S-cKO mice, consistent with our previous research. Further, expression levels of Sox2 declined in the RGL-derived cells in the SVZ of GD3S-cKO mice (0.583 ± 0.062-fold vs. control) (Figure 4g). The Nestin and GFAP double-positive RGL population were reduced in the SVZ of the GD3S-cKO mouse brain at 1 month after tamoxifen injection (Figure 3c). Nestin+ GFAP+ RGLs have been considered to be largely quiescent NSCs residing in the postnatal brain. Taken, together, these results indicate that GD3 in the RGLs is required to maintain the NSC pool in the SVZ.

### 3.3 Reduction of RGL-derived neuronal cells in the RMS-OB results in impairment of olfaction in GD3S-cKO mice

For neurogenesis in the OB, the immature neurons migrate through the RMS after being produced in the SVZ (Belvindrah et al., 2009; Kaneko et al., 2017; Lazarini & Lledo, 2011). SVZ-derived NPCs take about two weeks to reach the OB through the RMS and become matured (Alonso et al., 2006; Hack et al., 2005; Zhao et al., 2008). DCX-expressing NPCs migrate through the RMS towards the OB (Akter et al., 2021; Bressan & Saghatelyan, 2020; Capilla-Gonzalez et al., 2015; Doetsch & Alvarez-Buylla, 1996; Kaneko et al., 2017). The shrunk pool of NSCs points to a decrease of DCX+ neuronal precursors in the RMS and newborn neurons in the OB. To confirm functional significance of GD3 on postnatal neurogenesis, we knocked out GD3S at 1-month-old and observed the RMS at 4-month-old (Figure 5a). We found that the DCX expression in RGL-derived cells was declined in the GFAP+ RMS zones (Figure 5b). NPCs in the RMS zones generate mature neurons into the GCL and the PGL of the OB (Defterali et al., 2021; Lois & Alvarez-Buylla, 1994; Takahashi et al., 2018). We found the widespread radial distribution of DCX-expressing NPCs in the GCL as reported (Belvindrah et al., 2011; Kim et al., 2011; Kohl et al., 2010), regardless of positive or negative for Tmt in the control mice (Figure 5c). However, Tmt+DCX+ cells were rarely found in the GCL of GD3S-cKO, as consistent with decreased NSCs and NPCs in the SVZ and the RMS, respectively (Figure 4,5b). Since the mature neurons in the PGL express NeuN (Maier et al., 2017; Panzanelli et al., 2007; Schweyer et al., 2019), we quantified the NeuN+ cells among the Tmt+ RGL-derived cells as mature newborn neurons in the PGL (Figure 5d). The number of mature newborn neuron (Tmt+NeuN+) in the PGL of GD3-cKO mice was about 40% lower compared with control group (control; 6.95 ± 0.952% vs. GD3S-cKO; 4.30 ± 0.421) (Figure 5e). The results indicated that a decrease in the NSC pool in the SVZ leads to fewer neurogenesis in the PGL of the OB in the GD3S-cKO mouse brain.

Neurogenesis in the OB is considered essential for the proper olfaction (Brill et al., 2009; Hack et al., 2005; Miyamichi et al., 2011; Mizrahi et al., 2006). Neurogenesis continues in the SVZ throughout life, and RGLs from the SVZ contribute to those OB neurons that are integrated in the existing neuronal circuit (Nagayama et al., 2014). It is estimated that 10,000 new neurons are generated in the SVZ of adult mice daily and are incorporated into the OB via the RMS. To investigate the possible contribution of downregulated neurogenesis in the GD3S-cKO mice to the ability to smell, we performed the buried pellet test (Figure 6a,b), in which mice are expected to find the buried food pellet under the bedding in a cage, allowing us to assess the olfactory performance by measuring the latency needed (Alberts & Galef, 1971; Bermúdez et al., 2019; Fleming et al., 2008). As expected, we found that GD3S-cKO mice needed twice longer latency to find the food pellet beneath the bedding, suggesting inferior olfactory performance compared to the control mice (control; 56.6 ± 10.8 sec vs. GD3S-cKO; 102.4 ± 15.1 sec) (Figure 6c). These results indicate that GD3 play crucial role in maintaining olfaction.

### 3.4 Roles of GD3 in NSCs of the dentate gyrus and memory function

The DG of the hippocampus is a neurogenic region in the mammalian adult brain (Bermúdez et al., 2019; Gonçalves et al., 2016; Obernier & Alvarez-Buylla, 2019). It has been implicated that newborn neurons are involved in hippocampal-dependent cognition and mood control (Abbott & Nigussie, 2020; Anacker & Hen, 2017). We have reported that GD3 is involved in the maintenance of the function of newborn neurons in the DG by investigating in global GD3S-KO mice (Tang et al., 2021). It is unclear whether loss of GD3 in embryonic stage or lack of GD3 function in postnatal NSCs contribute to cognitive impairment. Since the Gli1 promoter is known to be activated in NSCs in the subgranular zone (SGZ) of the DG (Bin Imtiaz & Jessberger, 2021), we analyzed the proliferating activity of NSCs in the DG after depleted GD3S by administration of tamoxifen (Figure 7a). The RGL-derived cells of GD3-cKO mice showed impaired the ability of dividing as the reduced PCNA expressing proliferative Tmt+Sox2+ cells in the SGZ of the DG (control; 12.3 ± 3.58% vs. GD3S-cKO; 1.62 ± 1.02%) (Figure 7b,c). Next, we investigated newborn neurons in the DG of the GD3-cKO mouse brain. Tmt+NeuN+ neurons, which had been produced after tamoxifen treatment, appeared to tended to be decreased in the DG of GD3S-cKO mice (control; 3.98±1.20 cells/mm^2^ vs. GD3S-cKO; 3.12±0.37 cells/mm^2^) (Figure 7d,e). These decreased trends of neurogenesis imply the impairment of hippocampal-dependent memory. Since the neurogenesis has been associated with contextual object recognition memory (Faivre et al., 2011; Jessberger et al., 2009; Taha et al., 2020; White et al., 2020), we performed a novel object recognition test in a context (Goulart et al., 2010; Leger et al., 2013; Meziane et al., 1998; Murai et al., 2007) to assess the relationship between postnatal GD3 and hippocampal-dependent learning and memory. The 10 min habituation without any object was followed by the 5 min training session with two identical objects to be explored. Then mice were subjected to a 5 min trial phase with a novel object for 1h and 24h after the training as a memory retention test (Figure 8b). Memory for the familiar object was evaluated by the ratio of the spent time exploring the novel object which includes contacting by sniffing, touching, and climbing, versus total exploration time. The exploration time for the two identical objects was very similar, showing no spatial bias in these mice (Figure 8c,d). As expected, control mice preferred exploring novel objects over familiar ones, possibly due to their instinctive preference for novelty, however, significantly less spent time with the novel object was observed in GD3S-cKO mice in 1h memory retention test (control; 60.6 ± 4.41% vs. GD3S-cKO; 27.6 ± 5.42%) (Figure 8e). Further, GD3S-cKO mice spent less time with a novel object in 24h retention test compared to the control (control; 75.3 ± 5.20% vs. GD3S-cKO; 55.5 ± 5.20%) (Figure 8f). These results indicate that GD3S-cKO mice carry deficits in contextual novel object recognition. Collectively, these results suggest that GD3 is involved in memory possibly by NSC maintenance and following neurogenesis in the DG.

## 4 DISCUSSION

In this study, we demonstrated that (1) postnatal GD3 is required to maintain the quiescent RGLs in the SVZ, (2) postnatal deletion of GD3 in the RGLs leads to impaired neurogenesis in the OB with coupled with a deficiency of the NSC pools in the SVZ, (3) the reduced neurogenesis in the GD3S-cKO brain causes olfactory dysfunction, (4) loss of GD3 in RGLs also results in the decreased NSC pools and neurogenesis in the DG, and (5) postnatal GD3 deletion in RGLs leads to impairment of memory functions. These results suggest that GD3 regulates NSC fate determination to maintain the postnatal NSC pool and the ability of NSCs to undergo adult neurogenesis.

Postnatal neurogenesis is governed by the balance between quiescent and activated RGLs in the SVZ and the DG. Different cell types that have distinct functional roles are present in a NSC niche. Type B cells are reversibly quiescent RGLs in the SVZ and express GFAP. The quiescent type B1 radial glial cells are morphologically and phenotypically distinguished from the activated cells (no contacting cilium). Quiescent NSCs express platelet-derived growth factor receptor beta (PDGFRβ), and loss of PDGFRβ was reported to release the NSCs from quiescence (Delgado et al., 2021). The functional activities of proteins, such as growth factor receptors, are highly dependent upon their molecular environments. Cells as well as their subcellular organelles are surrounded by biological lipid membranes that define their individual cellular shape, help maintain cellular organization, and regulate signal transduction. Intriguingly, postnatal GD3-deficient RGLs have less radial glia-like morphology compared with control RGLs (Figure 2b). The cilium formation in RGLs is considered to have important functions to transduce the ventricular signals for the regulation of postnatal neurogenesis (Delgado et al., 2021; Falcao et al., 2012; Lehtinen et al., 2011; Tong et al., 2014). GD3 may contribute to radial glia-like morphological characteristics with cilium formation to receive the signals from cerebrospinal fluid for sustaining stemness of RGLs.

GD3 is a major glycolipid specie in developing and immature neuroectodermal populations in the rodent brain (Goldman et al., 1984). GD3 is predominant species (>80%) in the embryonic and adult NSCs and GD3 has been considered an NSC marker (Nakatani et al., 2010). In the adult stage, GD3 expression is limited to the neurogenic niches, including the SVZ and the DG, in the mouse brain. Global GD3S-KO mice show decreased the NSC pools, impaired postnatal neurogenesis, and decreased granule cells were observed in the OB and the DG (Itokazu et al., 2018; Wang et al., 2014; Wang & Yu, 2013). Our current study presents that RGL-specific inducible GD3 deletion promotes RGL activation state and reduced the NSC pool followed by declined neurogenesis in the OB and the DG of the postnatal mouse brain. The decreased neurogenesis in GD3S-cKO mice may lead to olfactory and memory dysfunctions. DCX+ immature neuronal cells in the RMS are suggested to be matured and integrated into neuronal circuits in the OB. Gli1+ RGL-derived neurons may contribute to olfaction, since GD3S-cKO mice showed longer latency in the buried pellet test, that is used to assess the ability to smell. Our observation of reduced Tmt+ cells in the GCL and periglomerular regions is in agreement with the previous reports that Gli1-expressing NSCs in the SVZ generate deep granule interneurons and a subpopulation of periglomerular neurons (Ihrie & Alvarez-Buylla, 2011). Hereafter, the subtypes of decreased neurons in the PGL in GD3S-cKO cells need to be identified since it would have a critical role in olfaction. DG-ablated rodent models reveals object recognition but failed to retain contextual-object memory recognition (Dees & Kesner, 2013). Although the DG-ablation models have shown that the DG is not indispensable for object recognition and its memory formation, it does not exclude the possibility that neurogenesis in the DG is associated with contextual object recognition and memory retention. In fact, functional association between neurogenesis and object memory retention in a context has been reported (Faivre et al., 2011; Jessberger et al., 2009; McGinley et al., 2018; Pardo et al., 2015; Taha et al., 2020; White et al., 2020). Moreover, Luna et al. has shown that adult-born granule cells in the DG contribute to the recognition of a novel object as contextual information in response to lateral entorhinal cortex (LEC) inputs (Luna et al., 2019). Neuronal network function between the DG and the LEC in the GD3S-cKO brain needs to be investigated. The morphological changes that occur as RGLs mature to neurons during neurogenesis requires a huge amount of energy (Son & Han, 2018). The mitochondrion, the main intracellular organelle for producing adenosine triphosphate (ATP), plays a crucial role in adult neurogenesis (Beckervordersandforth et al., 2017; Son & Han, 2018). We previously discovered that a mitochondrial fission protein, the dynamin-related protein-1 (Drp1), as a GD3-binding protein, and GD3 regulates mitochondrial dynamics. In global GD3S-KO mice, GD3S-KO adult-born neurons exhibited abnormal shorter mitochondria with a smaller aspect ratio, suggesting that mitochondria were fragmented (Tang et al., 2021). This mitochondrial fragmentation might occur in the GD3S-cKO mice. In accordance with those GD3 functions, GD3 could give a potential therapeutic strategy for the multiple symptoms caused by the declined the NSC pools and followed by neurogenesis.

The biological significance of adult neurogenesis in humans is still under debate (Boldrini et al., 2018; Kempermann et al., 2018; Moreno-Jimenez et al., 2019; Paredes et al., 2018; Sorrells et al., 2018; Tobin et al., 2019), however, the detection of adult neurogenesis has suggested that the adult brain exhibits more plasticity than previously thought, and that this contributes to memory and the pathogenesis of neurodegenerative diseases (Berger et al., 2020; Boldrini et al., 2018; Gage, 2004, 2019; Moreno-Jimenez et al., 2019; Tobin et al., 2019). Recently, we reported that exogenous administration of GD3 significantly restored the NSC pools and enhanced the stemness of NSCs with multipotency and self-renewal in the A53T a Synuclein-overexpressing Parkinson’s disease model (PD mouse), 5XFAD transgenic, and global GD3S-KO mice (Fuchigami et al., 2023; Itokazu et al., 2019). Our most recent study has shown that intranasal administration of GD3 recovered adult neurogenesis and olfaction in 7-9-month-old PD and global GD3S-KO mice with an easy noninvasive method. As adult neurogenesis is suggested to contribute to a healthy life, intranasal GD3 supplement may amplify the numbers of stem cells in the SVZ and hippocampus in human to help replenish brain circuits with neurons and improve the olfaction and memory functions with neurodegenerative and psychiatric diseases.

We have shown that GD3 has an important role for providing functional platforms for specific molecular interactions at GD3 microdomains in biological membranes of NSCs. In this study, we demonstrated that GD3 sustains quiescent RGLs to preserve NSC pools and controls postnatal neurogenesis. Cell surface and intracellular GD3 microdomains will provide additional clues underlying the adult neurogenesis, which may be useful in developing novel strategies for disease treatment and neuronal regeneration.

## ACKNOWLEDGMENT

This work was supported by a National Institute of Neurological Disorders and Stroke grant (R01 NS100839 to R.K.Y. and Y.I.). We thank Dr. Dongpei Li, Dr. Fu-Lei Tang, Dr. Jing Wang, Dr. Yong Li, Dr. Quanuang Zhang Dr. MB Khan and Dr. David Hess for excellent technical support. The authors also wish to acknowledge the excellent infrastructural support of the Department of Neuroscience and Regenerative Medicine (Chair, Dr. Xin-Yun Lu), Medical College of Georgia at Augusta University.

## ETHICS APPOLOVAL

All animal experiments were approved by the Institutional Animal Care and Use Committee (IACUC) at Augusta University (AU) according to the National Institutes of Health (NIH) guidelines and performed with the approved animal protocols (references AUP 2009-0240 and 2014-0694).

